# Ubiquitin Ligase MARCH5 Regulates the Formation of Mitochondria Derived Pre-peroxisomes

**DOI:** 10.1101/2023.10.26.564169

**Authors:** Jun Zheng, Zhihe Cao, Jinhui Wang, Yusong Guo, Min Zhuang

## Abstract

Peroxisome biogenesis involves two pathways: growth and division from pre-existing mature peroxisomes and *de novo* biogenesis from the endoplasmic reticulum, with a contribution from mitochondria, particularly in peroxisome deficient cells. However, the key components that regulate peroxisome *de novo* biogenesis are largely unknown. Dual organelle localized ubiquitin ligase MARCH5 functions on peroxisomes to regulate pexophagy. Here, we show that mitochondria localized MARCH5 is essential for the formation of vesicles in the *de novo* biogenesis of peroxisomes from mitochondria. Loss of MARCH5 specifically impedes the budding of PEX3-containing vesicles from mitochondria, thereby blocking the formation of pre-peroxisomes. Overall, our study highlights a novel function of MARCH5 for mitochondria derived pre-peroxisomes, emphasizing MARCH5 as one key regulator to maintain peroxisome homeostasis.

## INTRODUCTION

Mitochondrial homeostasis is maintained by multiple cellular pathways. Misfolded outer membrane proteins are recognized and eliminated through the ubiquitin-proteasome system by E3 ubiquitin ligase^1^. Damaged mitochondria undergo complete encapsulation within autophagosomes and subsequent degradation in lysosomes^2^. Additionally, mitochondria-derived vesicles (MDVs) selectively bud off to remove partially damaged mitochondria^3^. Under oxidative stress, over oxidized proteins aggregate within the mitochondrial matrix, which in turn triggers vesicle formation^4^. A recent study has demonstrated that TOMM20^+^ MDVs play a crucial role in clearing fully assembled protein complexes such as TOM import complexes^5^. Therefore, MDVs are considered as an alternative pathway for mitochondrial quality control.

However, lysosomes are not the sole destination for MDVs. It has been reported that MDVs containing MAPL specifically target peroxisomes, and their formation is independent of the mitochondrial fission factor DRP1^6^. These findings links MDVs with peroxisomes together and suggest distinct mechanisms governing the formation of different MDVs.

Peroxisome biogenesis consists of two distinct pathways: the growth and division of mature peroxisomes, and the *de novo* biogenesis from the endoplasmic reticulum (ER)^7, 8^. These two pathways are conserved in both yeast and humans. In addition, mitochondria may contribute to the *de novo* biogenesis of peroxisomes under certain circumstances. For example, in PBD (peroxisome biogenesis disorder) patient-derived fibroblast, the newly synthesized peroxisomes are a hybrid of PEX16-mediated vesicles from the ER and PEX3-mediated vesicles from mitochondria^9^. This discovery emphasizes the crucial role of MDVs in the *de novo* biogenesis of peroxisomes in higher eukaryotes.

The E3 ubiquitin ligase MARCH5 plays crucial role in maintaining mitochondrial homeostasis by regulating the dynamics of mitochondria through ubiquitination of key proteins involved in fission and fusion, such as Drp1, Fis1 and MFN1/2^10–13^.

Disruption of MARCH5-mediated mitochondrial dynamics can result in cellular senescence or induce cell death^14, 15^. Furthermore, MARCH5 is also implicated in protein quality control. For example, it specifically recognizes and binds to mutated superoxide dismutase 1 and expanded polyglutamine aggregates that accumulate within mitochondria^16, 17^. It also facilitates the recruitment of cytosolic Parkin to impaired mitochondria during mitophagy and regulates mitophagy receptor FUNDC1 in hypoxia^18, 19^. More, it was reported MARCH5 is involved in mitochondrial translocase to counteract protein import^1^. Additionally, MARCH5 regulates ER-Mitochondria contacts^20^. Therefore, MARCH5 is involved in mitochondria dynamic, protein quality control and organelle contacting site.

We have previously shown that MARCH5 also localizes to peroxisomes to mediate pexophagy under mTOR inhibition^21^. We found MARCH5 interacts with peroxisomal proteins. Notably, in peroxisome deficient cells, some peroxisomal proteins, including PEX3, are enriched on mitochondria^9^. In this study, we address the role of mitochondria-localized MARCH5 in peroxisome function. More specifically, we found MARCH5 is essential for the formation of vesicles budding from mitochondria in peroxisome *de novo* biogenesis. Loss of enzymatic activity or C terminal domain of MARCH5 inhibits the formation of newly synthesized peroxisomes. Moreover, we found mitochondria localized PEX3 can actively bud to form MDVs and this process relied on the core domain of PEX3. Our results suggest indispensable role of MARCH5 and PEX3 for the formation of mitochondria derived pre-peroxisomes.

## RESULTS

### Establishing peroxisome *de novo* biogenesis in peroxisome deficient cells

To understand the process of peroxisome *de novo* biogenesis from mitochondria in peroxisome deficient cells, we generated a HeLa cell line deficient in peroxisomes by knocking out the endogenous PEX3 gene (PEX3 KO) (**Fig S1**). PEX3 is an essential protein responsible for localizing peroxisomal membrane proteins (PMPs). Deletion of PEX3 leads to mislocalization of PMPs and ultimately results in the absence of functional peroxisomes. The cytosolic distribution analysis of GFP-SKL (GFP fused with a peroxisome targeting sequence Serine-Lysine-Leucine) confirmed the lack of functional peroxisomes in the PEX3 KO cells (**Fig S1**).

Next, similar to previously described^9^, we characterized different stages of peroxisome *de novo* biogenesis by reintroducing YFP-fused PEX3 based on its cellular localization pattern. Initially, tubular structures were observed in the presence of PEX3 (stage 0), which gradually transformed into vesicle formation (stage I). Finally, punctate structures indicating matured functional peroxisome formation were observed (stage II) (**Fig 1A**). The transition from stage 0 to stage II was also evident between 24 hours and 72 hours after reintroduction of PEX3-YFP expression (**Fig 1B**).

**Figure 1.**
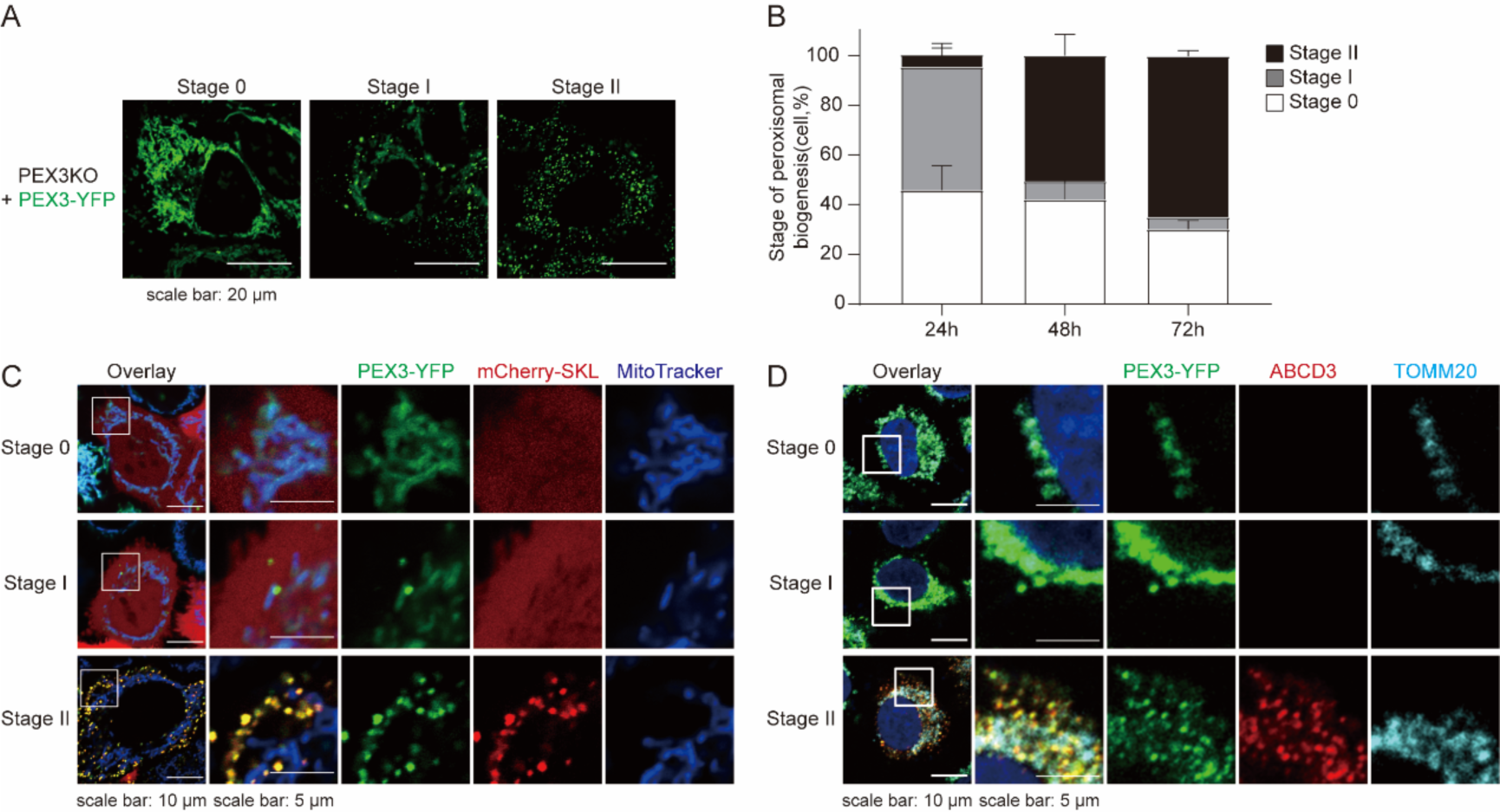
PEX3-YFP restore peroxisome *de novo* biogenesis in peroxisome deficient cell. (A) Representative image of PEX3 KO cells infected with PEX3-YFP. (B) The graph represents the percentage of cells at each stage of *de novo* biogenesis defined in a. Data from three biological replicates. Calculated cells were more than 100. (C) Representative image of mCherry-SKL stable expressed PEX3 KO cells infected with PEX3-YFP in each stage defined in (A). Cells were stained with MitoTracker Deep Red before imaging. (D) Representative image of PEX3 KO cells infected with PEX3-YFP in each stage. Cells were immnuostained with ABCD3 and TOMM20.

The subcellular localization of PEX3 during this process was also examined in the context of mitochondria. We stained mitochondria using MitoTracker or anti-TOMM20 antibody. The reintroduced PEX3-YFP was observed to specifically localize on the mitochondria (stage 0). Subsequently, it gradually formed pre-peroxisomes that exhibited negative staining for both peroxisomal membrane protein ABCD3 and peroxisomal matrix-targeted protein mCherry-SKL (stage I) before maturing into fully functional peroxisomes (stage II) (**Fig1C and 1D**).

Therefore, we reinstated peroxisome *de novo* biogenesis within mitochondria through exogenous expression of PEX3 in PEX3 knockout cells. This aligns with previous reports of *de novo* generation of peroxisomes by expressing PEX3 in patient fibroblasts lacking functional PEX3^9^.

### MARCH5 is required for peroxisome *de novo* biogenesis from mitochondria

To investigate the involvement of MARCH5 in peroxisome *de novo* biogenesis, we generated a double knockout cell line for both MARCH5 and PEX3 (MARCH5&PEX3 DKO, referred as DKO) (**Fig S2A**) and compared the biogenesis of peroxisomes in PEX3 KO and DKO cells. Using ABCD3 as the marker, reintroduction of PEX3-YFP restore the peroxisomes in PEX3 KO cells as expected, indicated by the co-localization of PEX3-YFP and ABCD3 outside of mitochondria. However, in the absence of MARCH5, both of PEX3-YFP and ABCD3 co-localize within mitochondria (**Fig 2A**), suggesting the absence of newly formed peroxisomes. This is further confirmed by using mCherry-SKL to detect mature peroxisomes (**Fig 2B**). In DKO cells, PEX3-YFP are retained on mitochondria and mCherry-SKL is evenly distributed in the cytosol. Statistical analysis further confirmed complete inhibition of peroxisome *de novo* biogenesis without the functional MARCH5 (**Fig 2C**). These findings collectively suggest that peroxisome *de novo* biogenesis was blocked in the absence of MARCH5.

**Figure 2.**
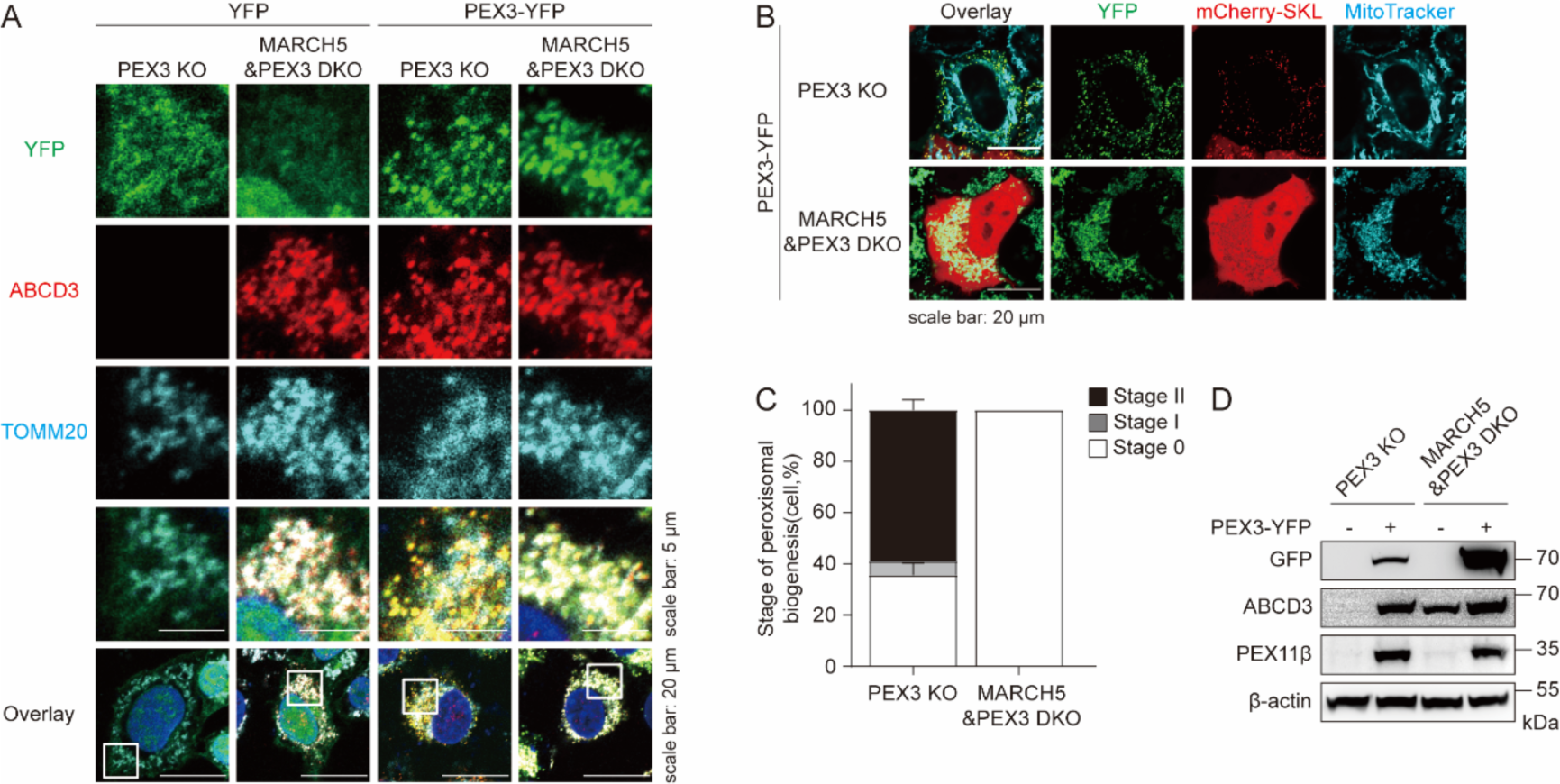
Loss of MARCH5 blocks peroxisome *de novo* biogenesis from mitochondria. **(A)** Representative image of PEX3 KO and MARCH5&PEX3 DKO cells infected with YFP or PEX3-YFP for 72 h. Cells were immunostained with ABCD3 and TOMM20 before imaging. **(B)** Representative image of mCherry-SKL stable expressed PEX3 KO and MARCH5&PEX3 DKO cells infected with PEX3-YFP for 72 h. Cells were stained with MitoTracker Deep Red before imaging. **(C)** The graph represents the percentage of cells at each stage of *de novo* biogenesis in a. Data from three biological replicates. Calculated cells were more than 100. **(D)** PEX3 KO and MARCH5&PEX3 DKO cells were infected by PEX3-YFP for 72 h. Expression of tagged and endogenous proteins is revealed by western blot, as indicated.

Unexpectedly, we observed the expression and localization of ABCD3 on mitochondria in DKO cells while ABCD3 cannot be detected in PEX3 KO cells (**Fig 2A**). We also examined ABCD3 expression levels in PEX3 knockout and MARCH5&PEX3 double knockout cells by western blots (**Fig 2D**). No ABCD3 can be detected in PEX3 KO cells, likely due to the lack of peroxisomes. The restoration of ABCD3 and another peroxisomal membrane protein PEX11β occurred upon the generation of newly formed peroxisomes. In DKO cells where MARCH5 is absent, we observed that ABCD3, but not PEX11β, exhibited endogenous expression regardless of exogenous PEX3 expression (**Fig 2D**). We previously reported MARCH5 as one ubiquitin ligase targeting ABCD3 for ubiquitination on peroxisomes^21^, it’s likely MARCH5 also mediated the degradation of ABCD3 on mitochondria.

### The RING domain and the C terminal peptide on MARCH5 are important for the ***de novo* biogenesis of peroxisomes**

The MARCH5 protein is composed of three distinct components: an N-terminal RING domain, a transmembrane domain consisting of four repeats, and a C-terminal peptide. The primary function of the RING domain is to facilitate the ubiquitination process for specific substrates^10–13^ . Mitochondrial localization relies on the transmembrane domain. Additionally, it has been reported that the C-terminal domain (CTD) plays a role in regulating peroxisome expansion^22^.

Subsequently, we employed different constructs of MARCH5 to investigate its functionality (**Fig S3**). The inhibition of *de novo* peroxisome biogenesis could be rescued by introducing wildtype MARCH5 into DKO cells but not by an enzymatically inactive form of MARCH5, indicating ubiquitination plays an important role in the formation of pre-peroxisomes. Additionally, swapping the RING domain of MARCH5 with RING domains from other ubiquitin ligases did not substitute for its function (**Fig 3A and 3B**), suggesting the RING domain of MARCH5 has its unique role to regulate peroxisome *de novo* biogenesis. Interestingly, we found that MARCH5 lacking the CTD did not rescue *de novo* peroxisome biogenesis either (**Fig 3C and 3D**). Taken together, these findings highlight the essential roles of both the RING domain and CTD in mediating MARCH5’s involvement in *de novo* peroxisome biogenesis.

**Figure 3.**
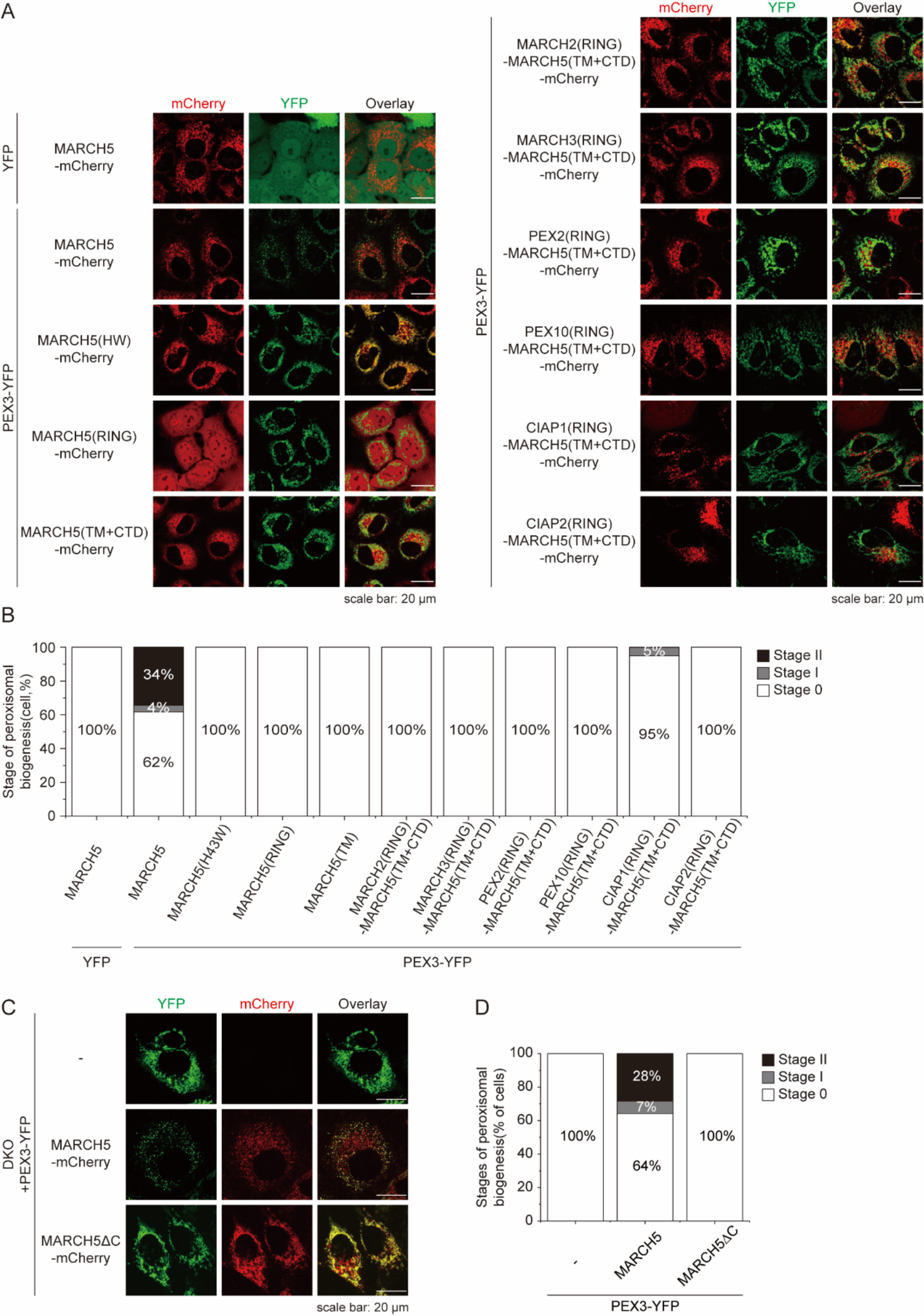
Both the RING and CTD domains are crucial for MARCH5-mediated pre-peroxisome formation. (A) mCherry fused wildtype MARCH5 or other variants were stable expressed in MARCH5&PEX3 DKO cells. Then, cells were infected with PEX3-YFP for 72 h and imaging. **(B)** The graph represents the percentage of cells at each stage of *de novo* biogenesis in A. Calculated cells were more than 100. **(C)** mCherry fused wildtype MARCH5 or MARCH5ΔC were stable expressed in MARCH5&PEX3 DKO cells. Then, cells were infected with PEX3-YFP for 72 h and imaging. **(D)** The graph represents the percentage of cells at each stage of *de novo* biogenesis in C. Calculated cells were more than 90.

### PEX3 can actively bud from mitochondria to form vesicles

The peroxisome deficient cells serve as a good model system to study the mitochondria derived vesicles (MDVs) since the pre-peroxisomes derived from mitochondria can be considered as a form of MDVs. In the absence of peroxisomes, multiple overexpressed peroxisomal proteins are targeted to mitochondria and may be susceptible to the mitochondrial quality control^23^.

To test whether PEX3 containing pre-peroxisomes under the surveillance by mitochondrial quality control, we treated the cells with autophagy inhibitor chloroquine (CQ). In addition to matured functional peroxisomes, we observed that a subset of PEX3-positive vesicles lacking the matrix protein mCherry-SKL in the presence of autophagy inhibitior CQ (**Fig 4A**), suggesting continuous degradation of these PEX3 containing vesicles under normal condition before they mature into functional peroxisomes. To further separate the budding step with the peroxisome maturation step, we examined whether PEX3 could bud from mitochondria in PEX19 knockout cells, where the peroxisomes are absent and the *de novo* biogenesis is also blocked due to the lack of PEX19 (**Fig 4B**). PEX19, a cytosolic protein chaperone involved in peroxisome membrane translocation along with PEX3, was completely knocked out resulting in the absence of cellular peroxisomes. We generated a PEX19 knockout cell line (PEX19 KO) and confirmed the absence of PEX19 through western blot (**Fig S4A**). PEX3-YFP localized mainly on mitochondria in PEX19 KO cells but a significant number of off-mitochondria PEX3-YFP vesicles appeared upon the treatment with CQ (**Fig 4C, Fig S4B**). This result suggests active budding of PEX3 from mitochondria to form pre-peroxisome-like vesicles, which are fast degraded if they cannot mature into functional peroxisomes.

**Figure 4.**
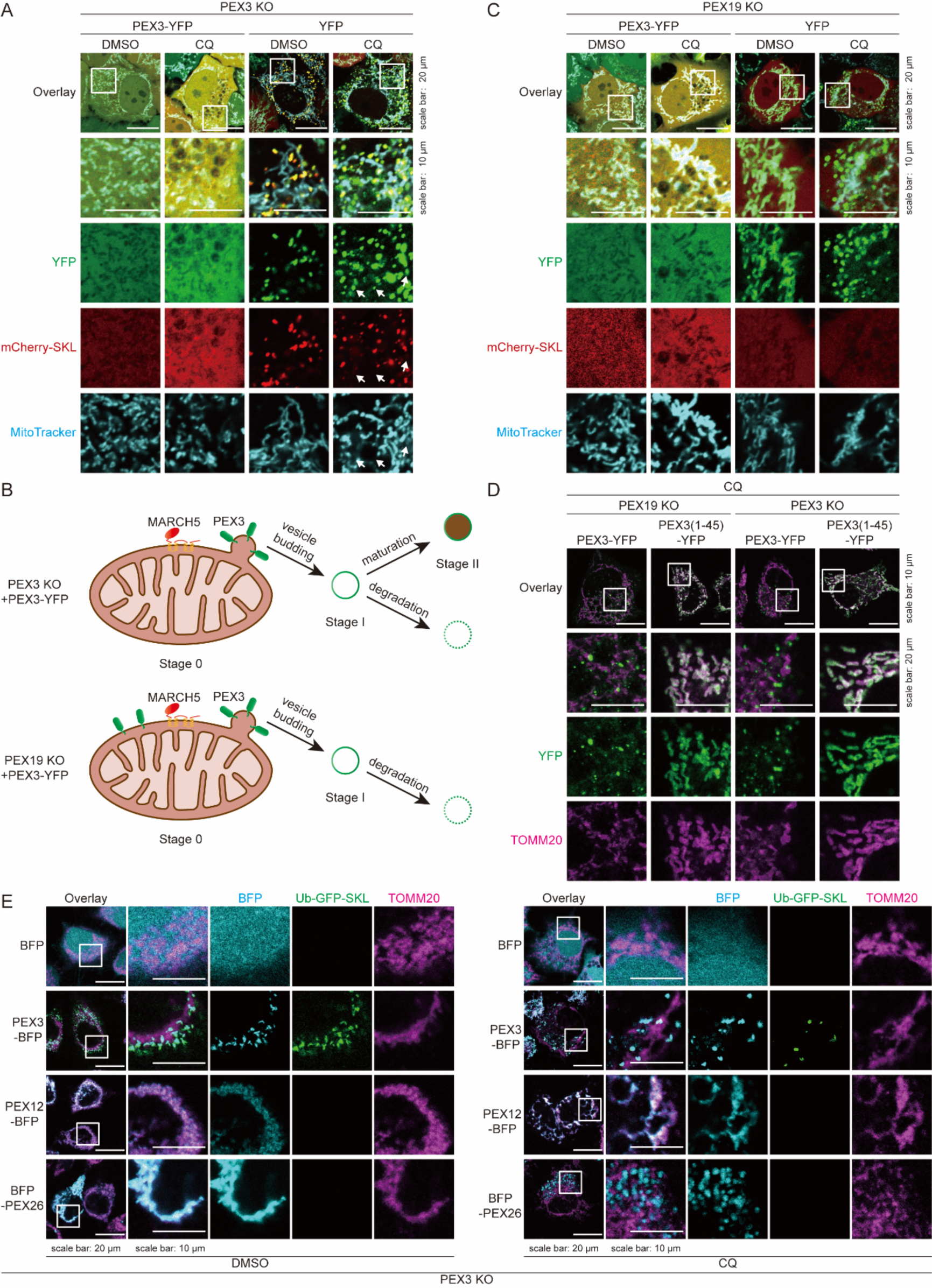
PEX3 can actively bud from mitochondria to form vesicles. (A) Representative image of mCherry-SKL stable expressed PEX3 KO cells infected with PEX3-YFP for 72 h. Cells were treated with DMSO or CQ (50 μM, 24 h), then stained with MitoTracker Deep Red before imaging. White arrows indicate unmatured pre-peroxisomes. **(B)** Schematic diagram of reintroducing PEX3 in PEX3 KO or PEX19 KO. **(C)** Representative image of mCherry-SKL stable expressed PEX19 KO cells infected with PEX3-YFP for 72 h. Cells were treated with DMSO or CQ (50 μM, 24 h), then stained with MitoTracker Deep Red before imaging. **(D)** Representative image of PEX3 KO or PEX19 KO cells infected with PEX3-YFP or PEX3(1-45)-YFP for 72 h. Cells were treated with CQ (50 μM, 24 h) and immunostained with TOMM20 before imaging. **(E)** Representative image of Ub-GFP-SKL stable expressed PEX3 KO cells infected with BFP fused PEX3 or other peroxisome membrane proteins for 72 h. Cells were treated with DMSO (left) or CQ (right) and immunostained with TOMM20 before imaging.

### Mitochondrial budding of PEX3-containing vesicles is dependent on the core domain of PEX3

We wonder if the packaging of PEX3 into the pre-peroxisomes is a passive or active procedure. PEX3 contains an N-terminal transmembrane helix and a C-terminal core domain. The N-terminal transmembrane helix (residues 1-45) of PEX3 is sufficient for its mitochondria localization (**Fig 4D**). PEX3 (1-45) mediates mitochondria localization of the fused YFP but cannot generate YFP-positive vesicles from the mitochondria, indicating that the C-terminal core domain of PEX3 is contributing to the vesicle budding. Despite being targeted to mitochondrial outer membrane by the same transmembrane domain, GFP and the counterpart, PEX3 core domain, contribute differently to the budding of vesicle structures, suggesting PEX3 actively induces pre-peroxisome formation.

PEX3-containing vesicles budded from mitochondria are different from those observed in yeast. A recent study revealed the formation of structures called mitochondria-derived compartments (MDCs) upon V-ATPase inhibition in yeast^24^. They also observed specific enrichment of TOMM70, an outer membrane protein of mitochondria, in these structures. Therefore, we examined whether PEX3-mediated vesicles are homologous to MDCs. Confocal imaging showed no significant colocalization between PEX3-positive vesicles and TOMM70 (**Fig S4C**). Furthermore, we found no overlap between PEX3-positive vesicles and LC3 (**Fig S4D**).

To investigate whether other peroxins induce the formation of MDVs, we expressed another two peroxisomal membrane proteins, PEX12 and PEX26 with fluorescent protein fusion. Both proteins are targeted to mitochondria in PEX3 knockout cells. In the presence of CQ, PEX26-containing vesicles can be observed but not with PEX12 (**Fig 4E**). Together, our data suggest whether the peroxisomal proteins induce the formation of degradable MDVs, are highly dependent on different peroxins.

### MARCH5 specifically regulates the budding of PEX3-containning vesicles from mitochondria

The introduction of PEX3 resulted in its retention on mitochondria in MARCH5&PEX3 DKO cells and no PEX3-containning vesicles can be observed in the presence of CQ (**Fig 5A**). However, it is not clear whether the absence of MARCH5 inhibits PEX3 budding or accelerates the degradation of budded vesicles to results in the major mitochondrial localization of PEX3. Since the budding of PEX3-containning vesicles and the maturation of these vesicles into peroxisomes can be de-coupled in PEX19 KO cells, it provides a system to investigate the role of MARCH5 in either PEX3 budding or pre-peroxisome maturation. Therefore, we generated a double knockout cell line for MARCH5 and PEX19 to examine the role of MARCH5 in MDV budding (**Fig S5A**). Overexpressed PEX3 formed vesicles upon treatment with CQ in PEX19 KO cells (**Fig 5B**), but was blocked on mitochondria in MARCH5&PEX19 DKO cells (**Fig 5B**), suggesting no PEX3-containing vesicles were formed in the absence of MARCH5. Given overexpressed PEX26 also forms MDVs that can be observed with CQ inhibition (**Fig 4D**), we examined the role of MARCH5 in generating PEX26-containing MDVs under the similar condition. However, the loss of MARCH5 did not affect the formation of vesicles containing PEX26 (**Fig 5C**).

**Figure 5.**
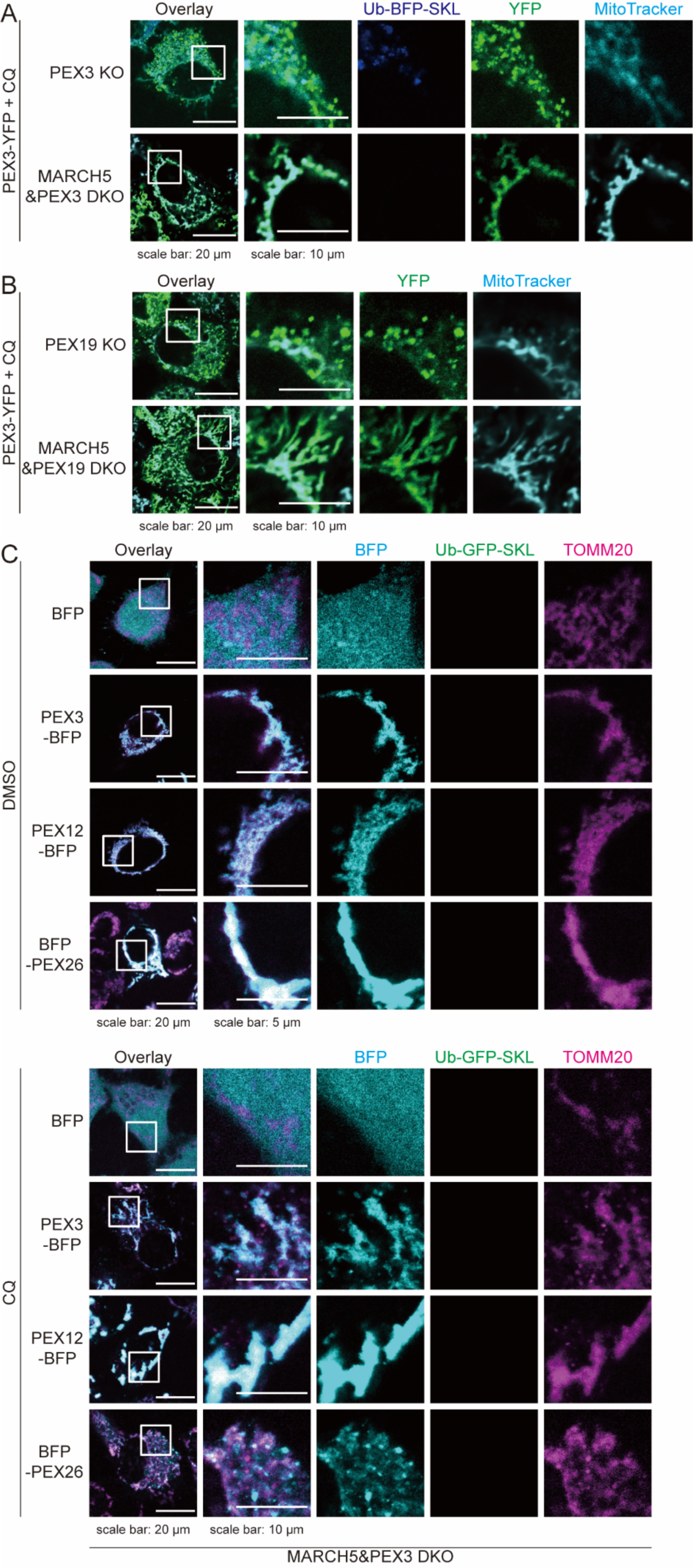
The loss of MARCH5 specifically inhibits the budding step of PEX3-containing vesicles. (A) Representative image of Ub-GFP-SKL stable expressed PEX3 KO or MARCH5&PEX3 DKO cells infected with PEX3-YFP for 72 h. Cells were treated with CQ (50 μM, 24 h) and stained with MitoTracker Deep Red before imaging. **(B)** Representative image of PEX19 KO or MARCH5&PEX19 DKO cells infected with PEX3-YFP for 72 h. Cells were treated with CQ (50 μM, 24 h) and stained with MitoTracker Deep Red before imaging. **(C)** Representative image of Ub-GFP-SKL stable expressed MARCH5&PEX3 DKO cells infected with BFP fused PEX3 or other peroxisome membrane proteins for 72 h. Cells were treated with DMSO (left) or CQ (right) and immunostained with TOMM20 before imaging.

**Figure 6.**
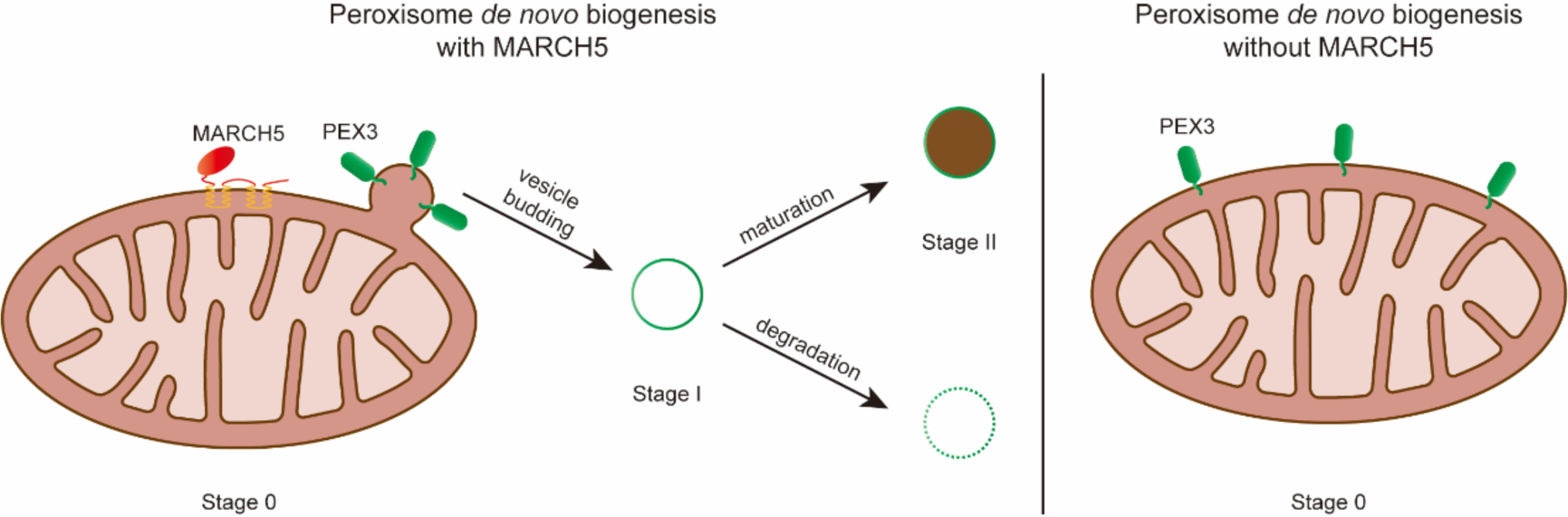
The working model for the role of MARCH5 in peroxisome *de novo* biogenesis. In peroxisome-deficient cells, reintroduced PEX3 localizes to the mitochondria and subsequently undergoes active budding to form vesicles that either mature into peroxisomes or are degraded. However, in the absence of MARCH5, PEX3 is unable to bud from the mitochondria and vesicle formation is blocked.

Overall, our findings suggest that MARCH5 specifically inhibits the formation of PEX3-positive vesicles derived from mitochondria, which is one of the major steps for the *de novo* biogenesis of peroxisomes.

## DISSCUSSION

The origin of organelles is a fundamental question in the field of cell biology. In yeast, peroxisomes are formed via distinct preperoxisomal vesicles arise from ER^25^. Recently, it has been proposed that the peroxisomes in human cells originates from vesicles derived from the ER and mitochondria^9^. The distinct origins between human and yeast emphasize the significance of mitochondria-derived vesicles in the *de novo* biogenesis of peroxisomes. However, the molecular mechanism underlying the formation of mitochondria-derived pre-peroxisomes remains largely unknown.

In this study, we generated a cellular model for peroxisome *de novo* biogenesis by introducing wild type PEX3 into PEX3 KO cells. This allows us to observe the newly synthesized peroxisomes derived from MDVs. We discovered that MARCH5 plays an indispensable role in the generation of pre-peroxisomes and the loss of MARCH5 ubiquitin ligase activity impedes the budding of PEX3-containing vesicles from mitochondria. In addition, the presence of PEX3 core domain is also necessary for the formation of pre-peroxisomes.

Are the PEX3-containing vesicles a special type of MDVs? MDVs can be categorized into two groups: those degraded by lysosomes and those mature or fuse to other organelles^26^. Although PEX3-containing vesicles can also be degraded, it has distinct proteins compositions from other MDVs. For instance, TOMM20, a recently identified MDV marker is not detected on PEX3-containing vesicles. Different MDV composition may indicate the different formation mechanism. DRP1 is required for TOMM20^+^ MDVs to exit from mitochondria^5^. Knockout of DRP1 and its receptors significantly reduces the number of TOMM20^+^ MDVs. However, the budding of PEX3-positive MDVs in peroxisome *de novo* biogenesis remains unaffected by the knockdown of DRP1^9^. Here we show MARCH5 specifically affects PEX3-positive MDVs but not the PEX26-positive MDVs, suggesting unique roles of both MARCH5 and PEX3 in the formation of pre-peroxisomes.

As a E3 ligase, MARCH5 functions via ubiquitinating a variety of substrates. For example, MARCH5 ubiquitinates several fission and fusion proteins to regulate mitochondria dynamics^10–13^. Given the classic mitochondria dynamic factors do not affect vesicles budding in peroxisome *de novo* biogenesis^9^, we speculate that there may be some unidentified proteins specifically responsible for this process, and their expression or ubiquitination level is regulated by MARCH5. Additionally, in our experiments, the RING domain of MARCH5 is necessary for peroxisome *de novo* biogenesis, and previous paper has reported the treatment of MG132 slow down this process^9^, we speculate the balance of ubiquitination of potential substrates is crucial to promote or inhibit peroxisome *de novo* biogenesis. Further research using genetic or molecular tools to identify new substrates of MARCH5 may shed light on this unexplored area.

### Key resources table

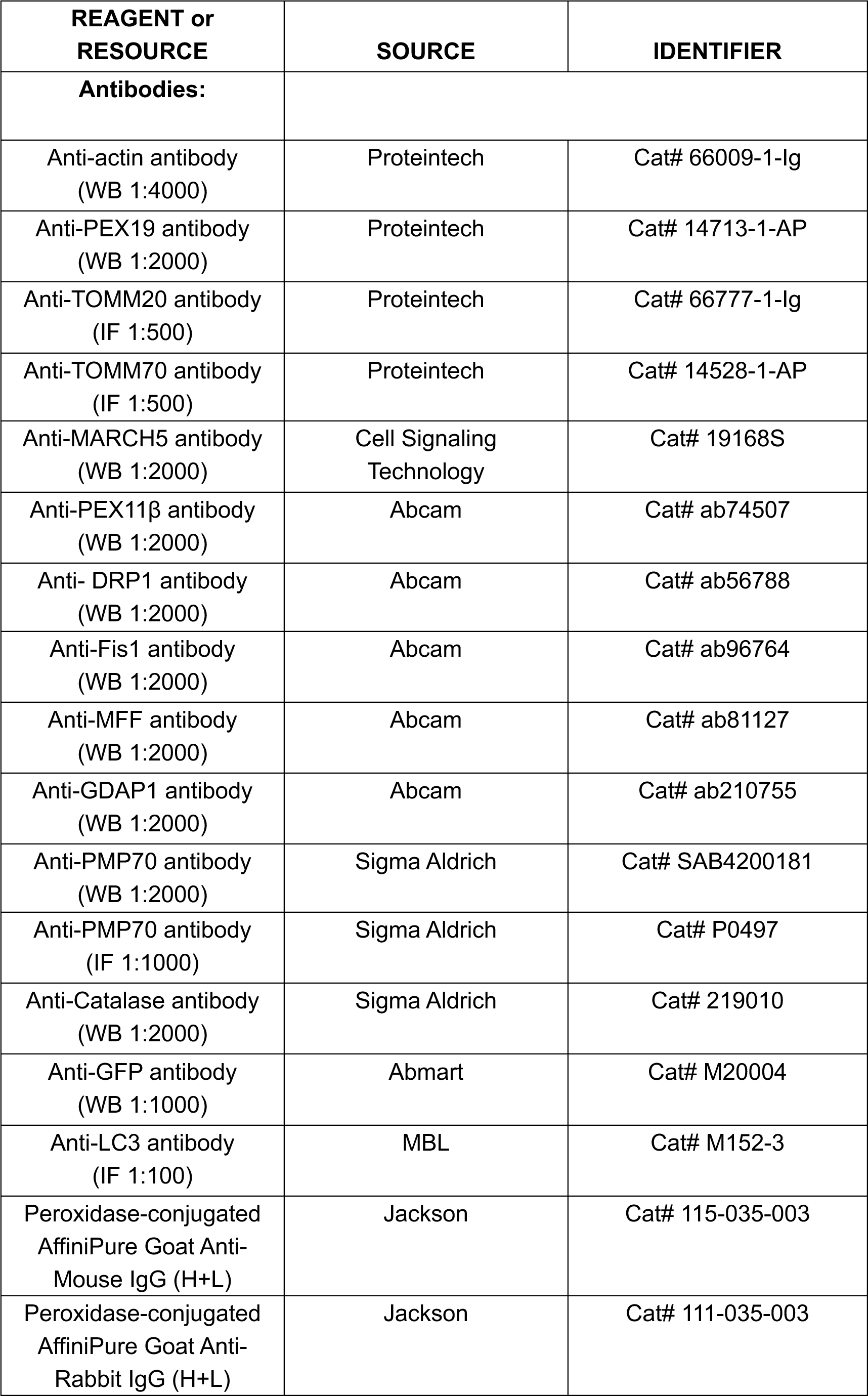
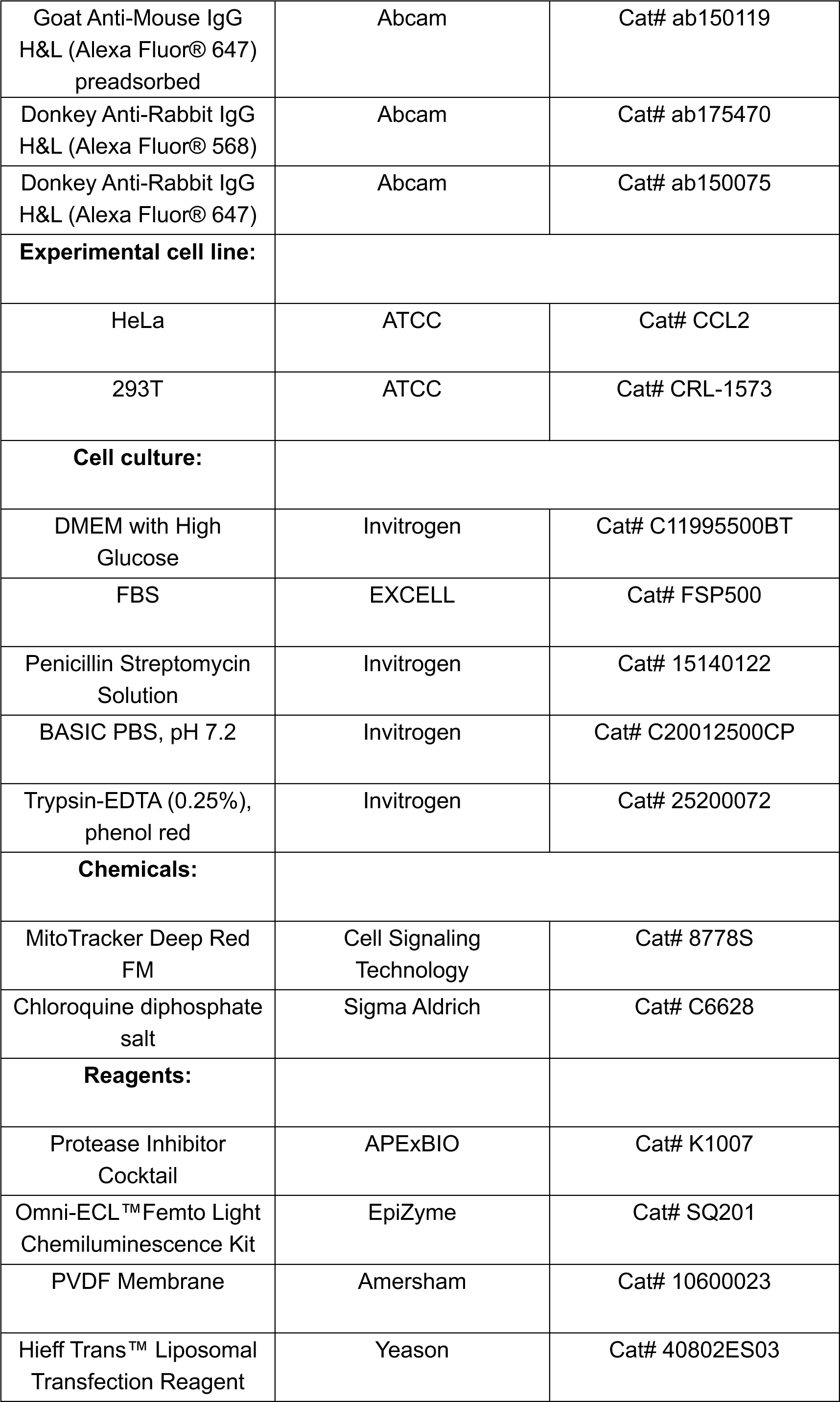
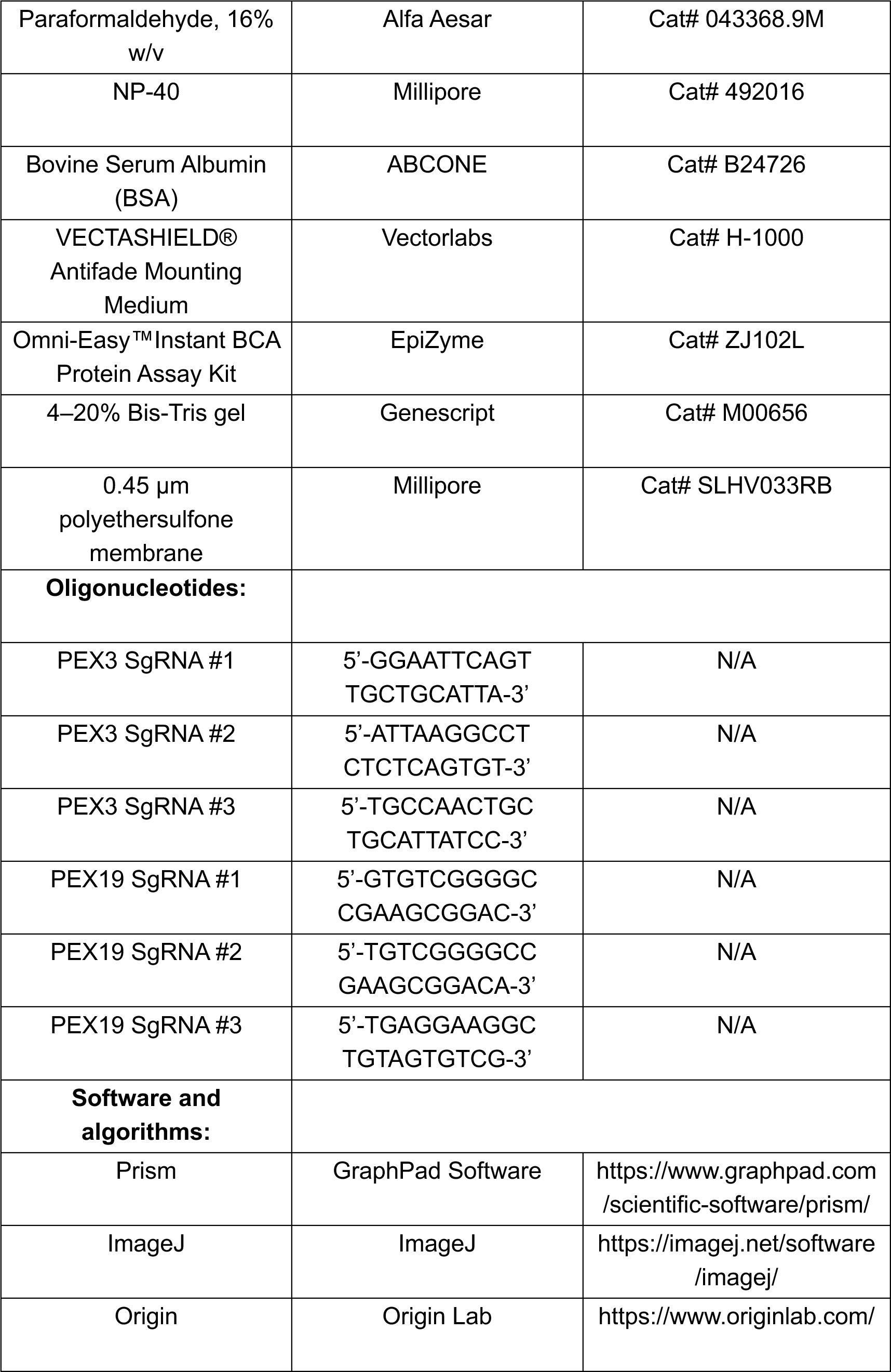

## METHODS

### Molecular cloning

We employed pHR_EF1a (derived from pHR_PGK, Addgene, #79120) for viral packaging. Fluorescent protein-fused PEX3 (Gene ID: 8504) and other PEX genes (PEX12 Gene ID: 5193; PEX26 Gene ID: 55670), MARCH5-mCherry and its variants (previous described^21^), mCherry-SKL, Ub-GFP-SKL, and Ub-BFP-SKL were integrated into pHR_EF1a via Gibson assembly utilizing BamHI and NotI restriction sites. All plasmids underwent Sanger sequencing to confirm their integrity.

### Lentivirus generation, infection and expression

Lentiviral vectors and their respective packaging vectors (pCMV-dR8.91 and pMD2G) were co-transfected into 293T cells at a molar ratio of 10:10:1, respectively. Following transfection, the media was changed to low-volume media (8 mL for a 10-cm dish) after 16 hours. The media was collected at 48 hours post-transfection, replaced with fresh media (8 mL), and collected again at 72 hours. To remove cell debris from the viral supernatant, centrifugation (10 min at 1,000 g) as well as filtration through a 0.45 µm polyethersulfone membrane were performed. For establishment of stable cell lines, cells were infected with the corresponding virus and sorted based on the expression of fluorescent proteins.

### Cell culture and transfection

293T (ATCC, CRL-1573) cells and HeLa (ATCC, CCL-2) were cultured in DMEM supplemented with 10% FBS in 5% CO2 at 37 °C. All media and FBS were obtained from Life Technologies and ExCell Bio. Transient transfections were performed using the Hieff Trans™ Liposomal Transfection Reagent according to the manufacturer’s instructions. All the cell lines were also tested and confirmed negative for mycoplasma.

### Immunofluorescence microscopy

The cells were plated on coverslips and incubated at 37 °C with 5% CO_2_ for 24 hours prior to staining. Subsequently, the cells were washed three times with 1× phosphate-buffered saline (PBS), fixed in 4% paraformaldehyde for 15 minutes, permeabilized with 0.1% NP-40 for 10 minutes, blocked with a solution of PBS containing 2.5% BSA for one hour at room temperature, and then incubated overnight at 4 °C with the primary antibody. Afterward, secondary antibodies were applied and allowed to react for one hour at a temperature of 37 °C. Following this step, DAPI staining was performed for two minutes and the samples were mounted using Mounting Medium. Confocal fluorescence imaging was conducted using a Nikon Ti-E+A1 R SI confocal microscope equipped with a high numerical aperture oil immersion objective lens (63× magnification; NA = 1.4) from Plan-Apochromat Lambda series along with an A1-DU4 detector unit utilizing photomultiplier tubes (PMT). Images were acquired as16-bit data files employing NIS-Elements C software. For visual presentation purposes, only brightness adjustments have been made.

### Generation of knockout cell lines by CRISPR/Cas9-based genome editing

HeLa cells lacking endogenous PEX3, PEX19 were generated by CRISPR/Cas9-mediated genome editing according to protocols published by the Zhang laboratory (rev20140509 at http://www.genome-engineering.org/crispr). MARCH5 knockout cell (MARCH5 KO) was established as described ^21^. MARCH5&PEX3 DKO and MARCH5&PEX19 DKO were generated based on MARCH5 KO and PEX19 KO, respectively. In brief, we used an online CRISPR design tool (http://crispr.mit.edu) to select individual guide RNA sequences targeting exon of genomic genes and designed the oligonucleotides. Oligonucleotides were phosphorylated, annealed and cloned into a modified pX330 (containing GFP) using BbsI restriction site. CRISPR constructs were first verified by sequencing and then transfected into cells. After 48 hours, transfected cells were diluted and plated for single clonal cell selections from GFP positive population. The GFP-positive cells were further screened for the absence of genes by immunoblotting using specific antibodies or by immunofluorescence imaging to verify the absence of peroxisomes.

### Western blotting assay

The cells were washed three times with PBS and subsequently lysed using a cell lysis buffer (composed of 50 mM Tris, 200 mM NaCl, 1% NP-40, pH 7.5) at a temperature of 4 °C for one hour. Prior to use, the lysis buffer was supplemented with a protease inhibitor cocktail at a concentration of 100×. The resulting lysates were then centrifuged at maximum speed (15,000g) and temperature of 4 °C for a duration of ten minutes. To determine protein levels, the supernatant was subjected to BCA Protein Assay. Subsequently, the cell lysates were separated on a 4–20% Bis-Tris gel, transferred onto PVDF membranes, and probed with antibodies.

### Quantification and statistical analysis

Quantitative data are presented as means ± standard deviation ^27^. The SD was calculated using Prism Graphpad, and error bars represent one SD from the mean.

---

**Figure S1.**
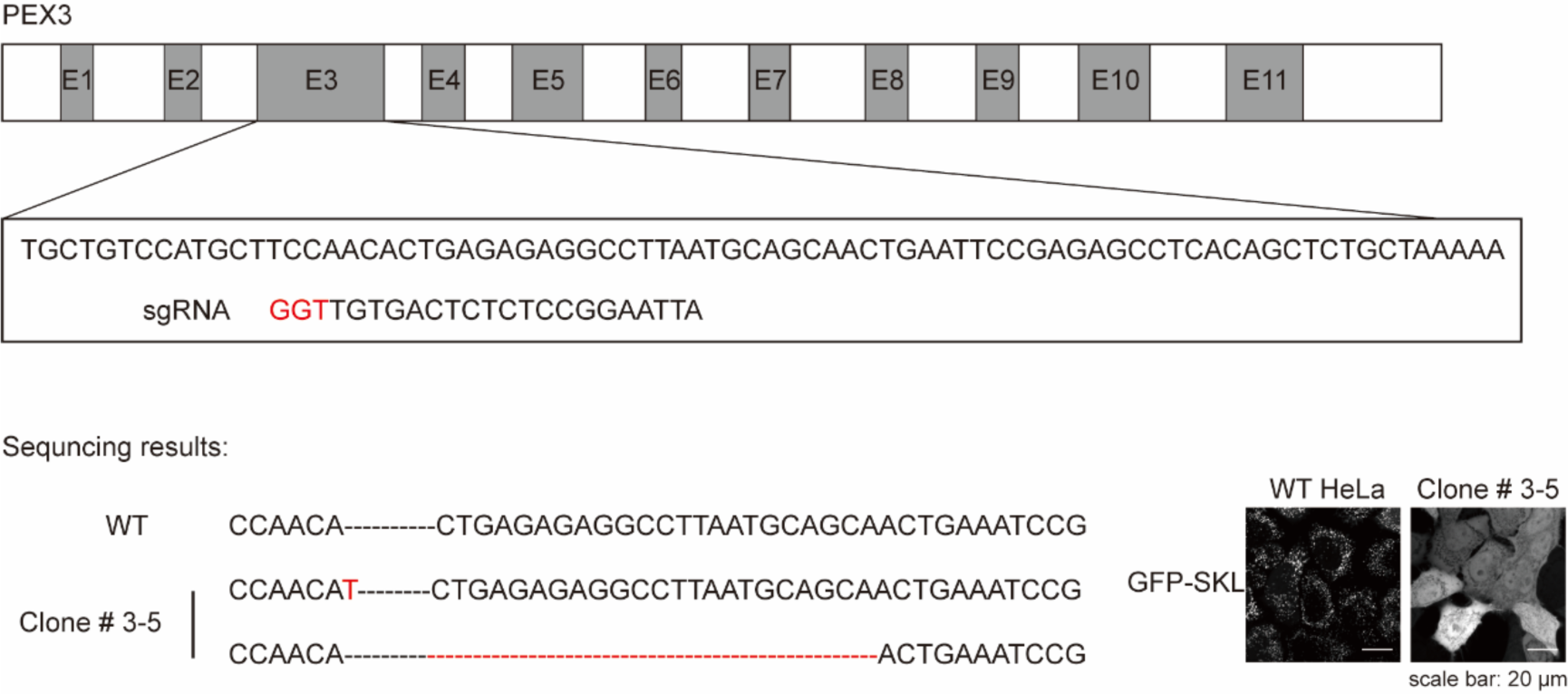
Generation of PEX3 KO cells. Schematic of the genome-editing strategy to knock out endogenous PEX3 in HeLa cells. Exons 1–11 (E1-E11) are indicated. Three designed sgRNAs were tested initially, and the validated sgRNA confirmed by sequencing, as shown. Protospacer adjacent motif sequences are depicted in red. The KO of PEX3 is confirmed by distribution of GFP-SKL.

**Figure S2.**
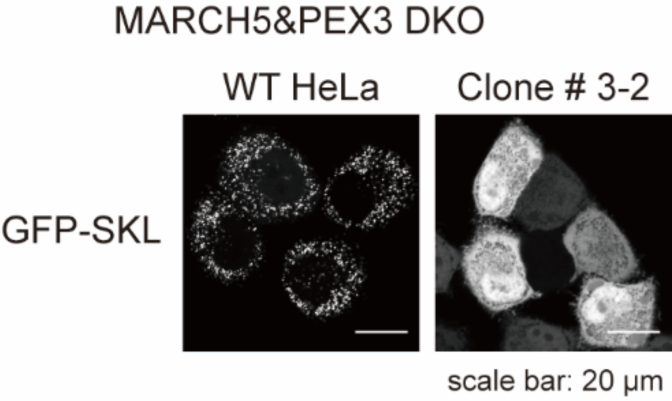
Validation of MARCH5&PEX3 DKO cells. MARCH5&PEX3 DKO cells were established based on MARCH5 KO. Endogenous PEX3 was knocked out using the same strategy in Figure S1. The KO of PEX3 is confirmed by distribution of GFP-SKL.

**Figure S3.**
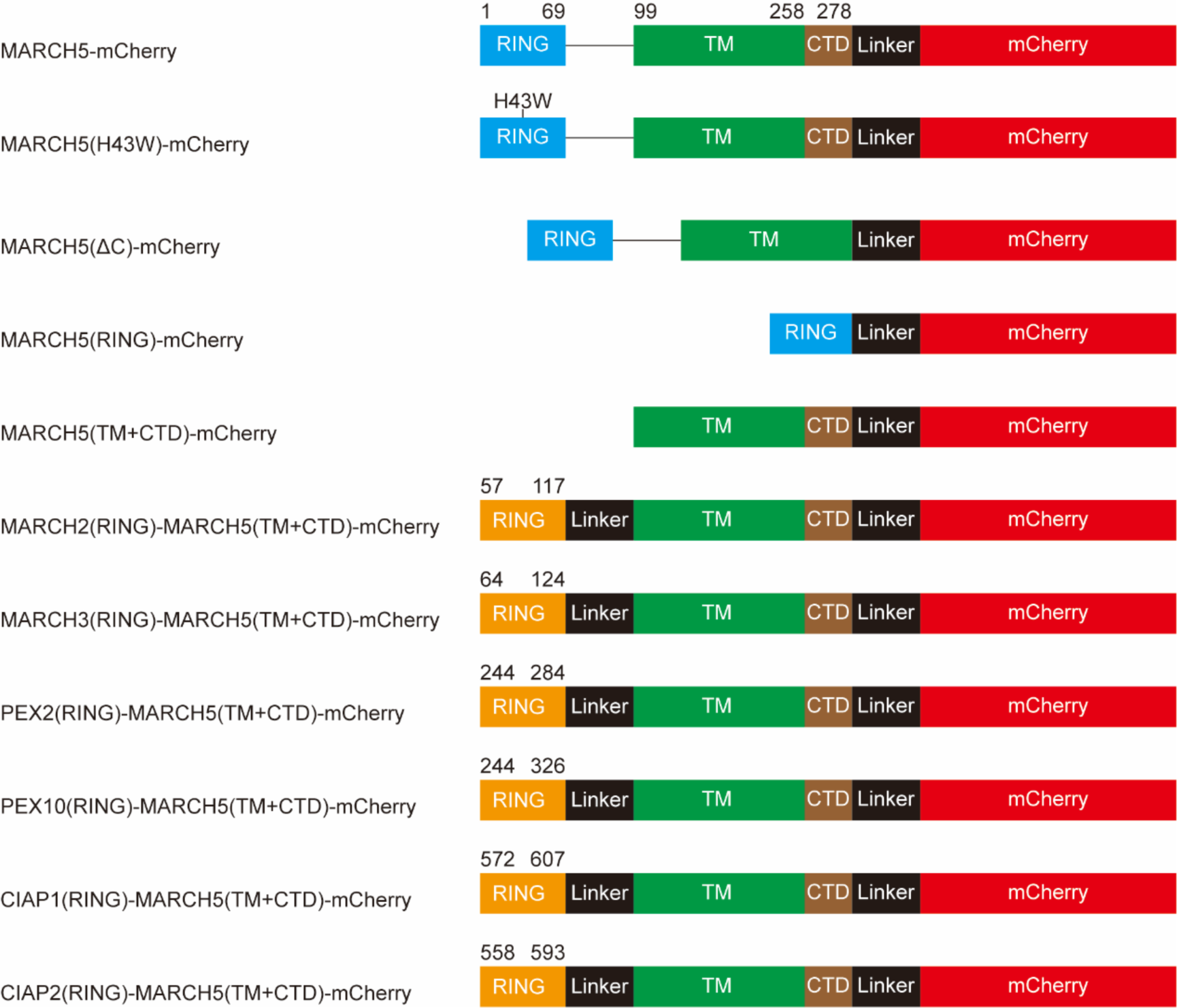
Constructs design of MARCH5 variants. TM, transmembrane domain. CTD, C terminal domain.

**Figure S4.**
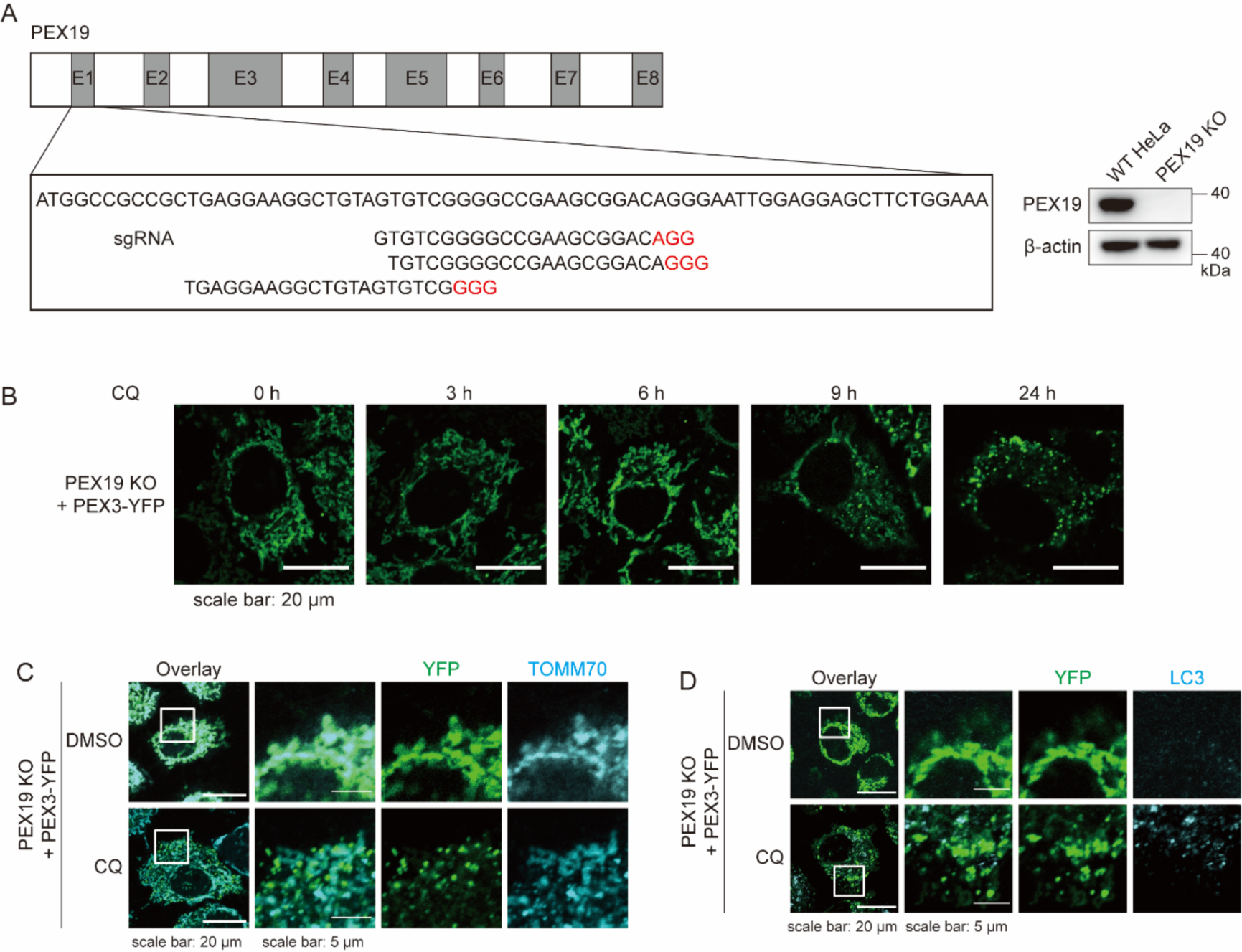
PEX3-positive MDVs in PEX19 KO cells. **(A)** Schematic of the genome-editing strategy to knock out endogenous PEX19 in HeLa cells. Exons 1–8 (E1-E8) are indicated. Three designed sgRNAs were tested initially as shown. Protospacer adjacent motif sequences are depicted in red. The KO of PEX19 is confirmed by WB. **(B)** Representative image of PEX3-YFP expressing in PEX19 KO cells treated with CQ for indicated time **(C-D)** Representative image of PEX3-YFP expressing in PEX19 KO cells. Cells were treated with CQ (50 μM, 24 h) and immune-stained with TOMM70**(C)** or LC3**(D)** before imaging

**Figure S5.**
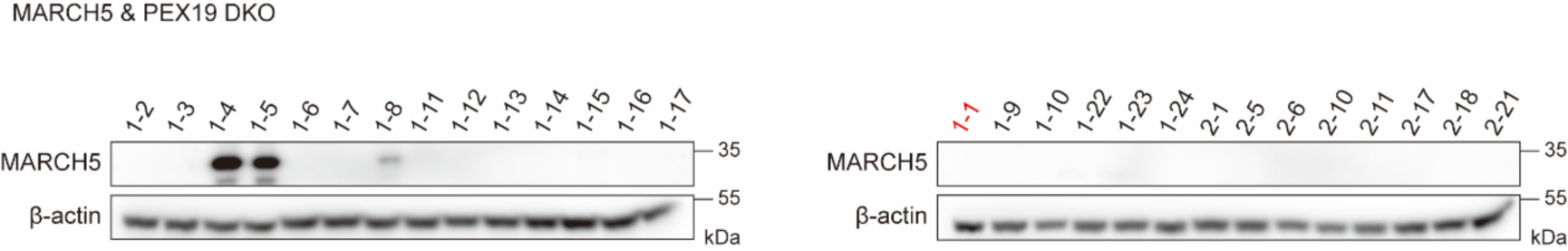
Generation of MARCH5&PEX19 DKO cells. MARCH5&PEX19 DKO cells were established based on PEX19 KO cells. The KO of MARCH5 is confirmed by WB. Clone #1-1 was used in this study.

## REFERENCE

1. Phu, L. et al. Dynamic Regulation of Mitochondrial Import by the Ubiquitin System. Molecular Cell 77, 1107–1123.e1110 (2020).

2. Youle, R.J. & Narendra, D.P. Mechanisms of mitophagy. Nature Reviews Molecular Cell Biology 12, 9–14 (2010).

3. McLelland, G.-L., Soubannier, V., Chen, C.X., McBride, H.M. & Fon, E.A. Parkin and PINK1 function in a vesicular trafficking pathway regulating mitochondrial quality control. The EMBO Journal 33, 282–295 (2014).

4. Shirihai, O.S. et al. Reconstitution of Mitochondria Derived Vesicle Formation Demonstrates Selective Enrichment of Oxidized Cargo. PLoS ONE 7 (2012).

5. König, T. et al. MIROs and DRP1 drive mitochondrial-derived vesicle biogenesis and promote quality control. Nature Cell Biology 23, 1271–1286 (2021).

6. Neuspiel, M. et al. Cargo-Selected Transport from the Mitochondria to Peroxisomes Is Mediated by Vesicular Carriers. Current Biology 18, 102–108 (2008).

7. Hoepfner, D., Schildknegt, D., Braakman, I., Philippsen, P. & Tabak, H.F. Contribution of the Endoplasmic Reticulum to Peroxisome Formation. Cell 122, 85–95 (2005).

8. Kragt, A., Voorn-Brouwer, T., van den Berg, M. & Distel, B. Endoplasmic Reticulum-directed Pex3p Routes to Peroxisomes and Restores Peroxisome Formation in a Saccharomyces cerevisiae pex3Δ Strain. Journal of Biological Chemistry 280, 34350–34357 (2005).

9. Sugiura, A., Mattie, S., Prudent, J. & McBride, H.M. Newly born peroxisomes are a hybrid of mitochondrial and ER-derived pre-peroxisomes. Nature 542, 251–254 (2017).

10. Karbowski, M., Neutzner, A. & Youle, R.J. The mitochondrial E3 ubiquitin ligase MARCH5 is required for Drp1 dependent mitochondrial division. The Journal of Cell Biology 178, 71–84 (2007).

11. Yonashiro, R. et al. A novel mitochondrial ubiquitin ligase plays a critical role in mitochondrial dynamics. The EMBO Journal 25, 3618–3626 (2006).

12. Park, Y.-Y. & Cho, H. Mitofusin 1 is degraded at G2/M phase through ubiquitylation by MARCH5. Cell Division 7, 25 (2012).

13. Kim, H.-J. et al. HDAC6 maintains mitochondrial connectivity under hypoxic stress by suppressing MARCH5/MITOL dependent MFN2 degradation. Biochemical and Biophysical Research Communications 464, 1235–1240 (2015).

14. Park, Y.-Y. et al. Loss of MARCH5 mitochondrial E3 ubiquitin ligase induces cellular senescence through dynamin-related protein 1 and mitofusin 1. Journal of Cell Science 123, 619–626 (2010).

15. Trounce, I.A. et al. Inactivation of MARCH5 Prevents Mitochondrial Fragmentation and Interferes with Cell Death in a Neuronal Cell Model. PLoS ONE 7 (2012).

16. Yonashiro, R. et al. Mitochondrial Ubiquitin Ligase MITOL Ubiquitinates Mutant SOD1 and Attenuates Mutant SOD1-induced Reactive Oxygen Species Generation. Molecular Biology of the Cell 20, 4524–4530 (2009).

17. Sugiura, A. et al. A mitochondrial ubiquitin ligase MITOL controls cell toxicity of polyglutamine-expanded protein. Mitochondrion 11, 139–146 (2011).

18. Koyano, F., Yamano, K., Kosako, H., Tanaka, K. & Matsuda, N. Parkin recruitment to impaired mitochondria for nonselective ubiquitylation is facilitated by MITOL. Journal of Biological Chemistry 294, 10300–10314 (2019).

19. Chen, Z. et al. Mitochondrial E3 ligase MARCH5 regulates FUNDC1 to fine-tune hypoxic mitophagy. EMBO reports 18, 495–509 (2017).

20. Sugiura, A. et al. MITOL Regulates Endoplasmic Reticulum-Mitochondria Contacts via Mitofusin2. Molecular Cell 51, 20–34 (2013).

21. Zheng, J., Chen, X., Liu, Q., Zhong, G. & Zhuang, M. Ubiquitin ligase MARCH5 localizes to peroxisomes to regulate pexophagy. Journal of Cell Biology 221 (2022).

22. Koyano, F. et al. Parkin-mediated ubiquitylation redistributes MITOL/March5 from mitochondria to peroxisomes. EMBO reports 20 (2019).

23. Nuebel, E. et al. The biochemical basis of mitochondrial dysfunction in Zellweger Spectrum Disorder. EMBO reports 22 (2021).

24. Schuler, M.-H. et al. Mitochondrial-derived compartments facilitate cellular adaptation to amino acid stress. Molecular Cell 81, 3786–3802.e3713 (2021).

25. van der Zand, A., Gent, J., Braakman, I. & Tabak, Henk F. Biochemically Distinct Vesicles from the Endoplasmic Reticulum Fuse to Form Peroxisomes. Cell 149, 397–409 (2012).

26. Sugiura, A., McLelland, G.L., Fon, E.A. & McBride, H.M. A new pathway for mitochondrial quality control: mitochondrial-derived vesicles. The EMBO Journal 33, 2142–2156 (2014).

27. Papsdorf, K. et al. Lipid droplets and peroxisomes are co-regulated to drive lifespan extension in response to mono-unsaturated fatty acids. Nat Cell Biol 25, 672–684 (2023).

